# MeCP2 controls dendritic morphogenesis via miR-199a-mediated Qki downregulation

**DOI:** 10.1101/2025.04.04.642981

**Authors:** Koichiro Irie, Hideyuki Nakashima, Masahiro Yamaguchi, Fumitaka Osakada, Norio Ozaki, Keita Tsujimura, Kinichi Nakashima

**Affiliations:** Department of Stem Cell Biology and Medicine, Graduate School of Medical Sciences, Kyushu University, 3-1-1 Maidashi, Higashi-ku, Fukuoka, Fukuoka 812-8582, Japan; Department of Neuroscience and Cell biology, Graduate School of Medicine, The University of Osaka, 2-2, Yamadaoka, Suita, Osaka, 565-0871, Japan; Laboratory of Cellular Pharmacology, Department of Basic Medicinal Sciences, Graduate School of Pharmaceutical Sciences, Nagoya University, Furo, Chikusa, Nagoya, Aichi, 464-8601, Japan; Department of Psychiatry, Graduate School of Medicine, Nagoya University, 65 Tsurumai, Showa-ku, Nagoya, Aichi, 466-8550, Japan; Research Unit for Developmental Disorders, Institute of Advanced Research, Nagoya University, Furo, Chikusa-ku, Nagoya, Aichi, 464-8601, Japan

## Abstract

Rett syndrome (RTT) (OMIM: 312750) is a severe neurodevelopmental disorder caused by mutations in the *MECP2* gene. Although decreased dendritic morphogenesis has been observed in the brain of RTT patients and mouse models, the molecular mechanisms underlying these dendritic anomalies remain unclear. We have previously shown that MeCP2 facilitates specific microRNA (miRNA) processing by associating with the miRNA microprocessor Drosha complex. In this study, we show that MeCP2 positively regulates dendritic formation via *miR-199a*, a specific target of the MeCP2-Drosha complex. Overexpression of MeCP2 and miR-199a promotes dendritic development such as increases in dendrite length, branching number, and complexity. In contrast, blocking miR-199a inhibited dendrite formation and abolished enhanced dendritic development induced by MeCP2 expression. We also demonstrate that the decreased dendrite outgrowth observed in MeCP2-deficient neurons could be rescued by miR-199a expression. In addition, we found that miR-199a targets the 3’ untranslated region of *quaking (Qki)*, a negative regulator of dendritic development, and downregulates its protein expression level. Furthermore, we report an increase in the Qki protein expression level in *miR-199a-2*-deficient brains and show that Qki knockdown restores the dendritic morphology of *miR-199a-2*-Knockout (KO) neurons. Taken together, these results suggest that the MeCP2/miR-199a/Qki axis is critical for proper dendritic development and its dysregulation contributes to the dendritic pathology in RTT.

## Introduction

Dendritic morphogenesis is critical for establishing functional neural circuits during brain development (1, 2), with dendrite anomalies associated with neurodevelopmental and neuropsychiatric disorders (3–5). Rett syndrome (RTT) is a progressive neurodevelopmental disorder caused by loss-of-function mutations in the X-linked *methyl-CpG binding protein 2 (MECP2)* gene (6). RTT patients develop normally until 6-18 months of age, beyond which there is an onset of regression and various neurological symptoms including seizures, stereotypies, loss of motor skills, autistic features, and mental retardation (7, 8). Studies on postmortem brains have revealed dendritic atrophy in RTT patients (9–11). Several mouse models lacking *Mecp2* (MeCP2-KO) or carrying a mutation on *Mecp2* (12–16) mimic many RTT phenotypes including motor impairments, seizures, and stereotypies (12, 13, 16). Reduced dendritic growth was observed in these models (17, 18) and dendritic atrophy was reported in cultured MeCP2-deficient neurons (19, 20). Therefore, impaired dendritic development may contribute to RTT pathophysiology, with MeCP2 regulating dendritic formation. However, little is known about the mechanisms underlying these phenomena.

MeCP2 belongs to the methylated DNA binding protein family and functions as a transcriptional repressor by recruiting co-repressors including histone deacetylases (HDACs) (21, 22), in addition to being a transcriptional activator and a splicing regulator (23, 24).

MicroRNAs (miRNAs) are small noncoding RNAs that regulate gene expression at the post-transcriptional level and are involved in many biological processes including cell proliferation, differentiation, and development (25, 26). Primary miRNAs (pri-miRNAs) are transcribed from DNA, processed into precursor miRNAs (pre-miRNAs) by the nucleus-associated Drosha complex, and transported to the cytoplasm by Exportin-5, where these are processed into mature miRNAs by Dicer. Mature miRNAs bind to the 3’ untranslated region (3’UTR) of target mRNAs and regulate gene expression by inhibiting their translation or degradation (27, 28). Increasing evidence suggests a key regulatory role of miRNAs in dendritic development. For example, miR-137 represses dendritic morphogenesis by inhibiting Mind bomb 1 expression (29) and miR-9 regulates dendritic formation by targeting the *repressor element-1 silencing transcription factor* (*REST*) gene (30). Recently, we have shown that miR-214 positively controls dendritic morphogenesis by regulating the expression of the schizophrenia-associated protein quaking (Qki) (31).

We have reported that MeCP2 facilitates specific pri-miRNA processing by associating with the Drosha complex and identified miR-199a as a target of the MeCP2-Drosha complex (32). We also showed that miR-199a-5p, a mature form of miR-199a, regulates mTOR signaling downstream of MeCP2 by inhibiting the expression of mTOR signal inhibitors, and dysregulation of the MeCP2/miR-199a-5p/mTOR signaling contributes to RTT neuronal phenotypes such as reduction in excitatory synaptic transmission and density and smaller neuronal soma size (32). Furthermore, we demonstrated that genetic deletion of miR-199a-2, a genomic loci of miR-199a, recapitulates RTT phenotypes in mice and leads to decreased brain mTOR activity (32). Although these findings identify miR-199a as a critical downstream target of MeCP2 in RTT pathology, its contribution to dendritic phenotype remains unexplored. Despite reports on suppression of dendritic development by MeCP2-mediated disruption of the Drosha-DGCR8 complex (33), this negative regulation of dendritic morphology by MeCP2 is inconsistent with the dendritic phenotype in postmortem brains of RTT individuals and animal models (9–11, 17, 18).

In this study, both miR-199a-5p and miR-199a-3p were found to contribute to dendrite development downstream of MeCP2. Overexpression and inhibition of miR-199a in primary cultures of hippocampal neurons promoted and impeded dendrite development, respectively. Importantly, blocking miR-199a-3p strongly inhibited dendritic morphogenesis and completely abolished MeCP2-induced enhancement of dendritic growth, highlighting the crucial role of miR-199a-3p in dendritic development. We also demonstrated that miR-199a overexpression rescued dendritic impairment in MeCP2-KO neurons. Additionally, miR-199a-3p was observed to target *Qki* mRNA and suppress Qki expression. Indeed, Qki expression was elevated in miR-199a-2-KO mice brain and *Qki* knockdown improved impaired dendritic development in miR-199a-2-KO neurons. These results suggest that the MeCP2/miR-199a/Qki pathway plays a critical role in dendritic development and its dysregulation accounts for the dendritic phenotype in RTT mouse models.

## Results

### Expression of MeCP2 correlates with mature-miR-199a production in developing hippocampal neurons

Although previous studies have shown that MeCP2 positively regulates neuronal maturation (34, 35), little is known about the mechanistic basis of its regulation of dendritic development. Previously, we have reported that MeCP2 promotes specific miRNA processing and identified miR-199a as a MeCP2 target accountable for the RTT phenotypes in mouse models of the disease (32). For this reason, we focused on the role of miR-199a for understanding MeCP2-regulated dendritic development of neurons. To explore the impact of the MeCP2/miR-199a pathway on dendritic morphogenesis, we examined the expression patterns of MeCP2 and miR-199a in cultures of developing hippocampal neurons. MeCP2 expression level gradually increased during neuronal development (Fig. 1A and B). The pri-miR-199a is expressed from 2 gene loci, *miR-199a-1* and *miR-199a-2*, and pre-miR-199a eventually generates 2 mature forms of miR-199a, i.e., miR-199a-5p and miR-199a-3p, which are highly conserved across mammalian species (Fig. 1C) (32). Expression levels of pri-miR-199a-1 and pri-miR-199a-2 in cultured hippocampal neurons were found to increase at an early stage and then gradually decrease at later stages of neuronal maturation (Fig. 1D and E), whereas that of the 2 types of mature miR-199a (miR-199a-5p and −3p) increased with neuronal dendrite growth (Fig. 1F and G). These results imply that the MeCP2/miR-199a axis has a role in the regulation of dendritic morphogenesis.

**Fig. 1.**
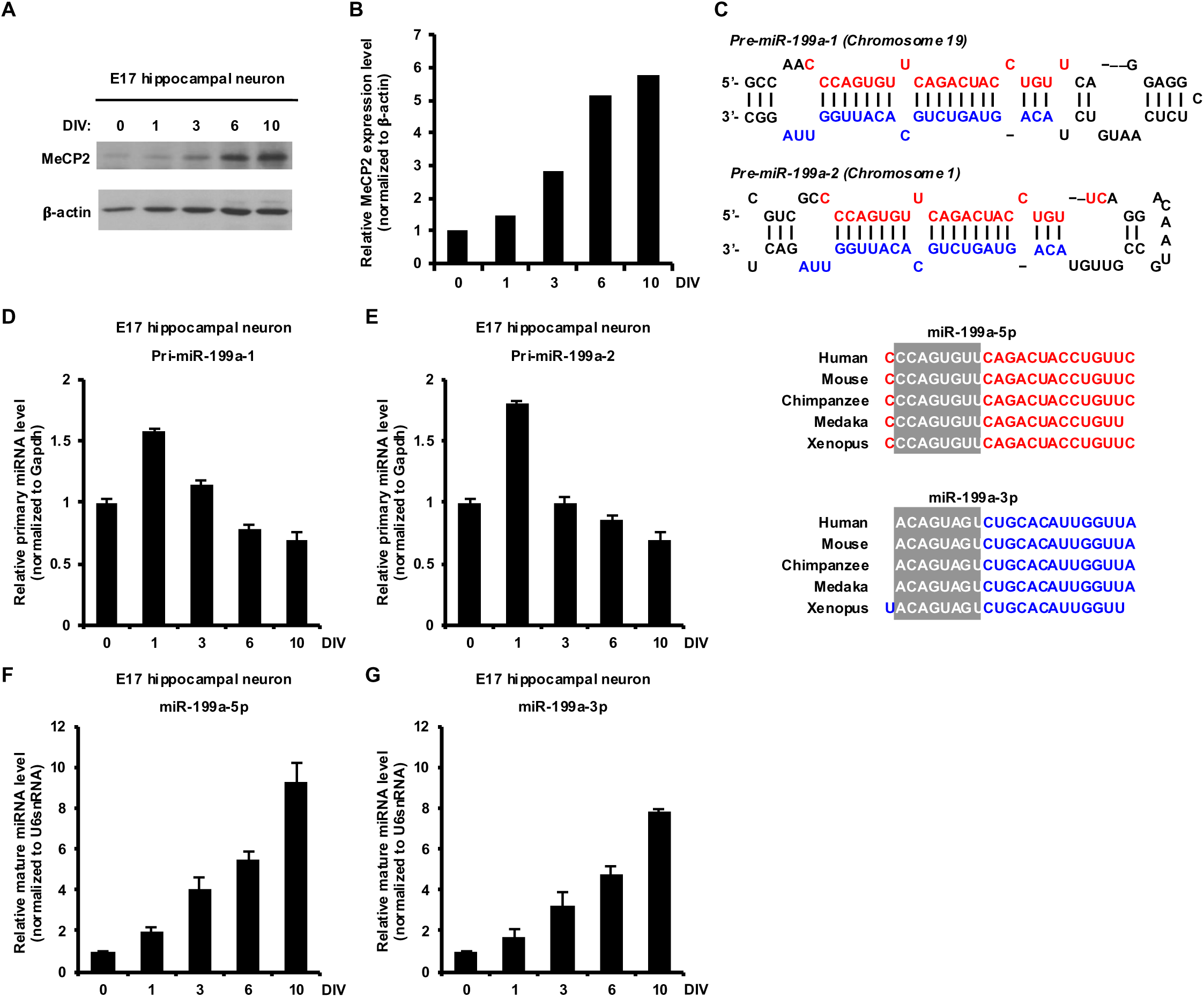
Expression patterns of MeCP2 and miR-199a in hippocampal neurons during development. *A*, Expression levels of MeCP2 were measured by immunoblotting in cultures of developing hippocampal neurons at 0, 1, 3, 6, and 10 days *in vitro* (DIV). *B*, Quantification of MeCP2 expression in *A*. *C*, Diagrammatic representation of the sequence of pre-miR-199a-1 and pre-miR-199a-2. The nucleotide sequences of mature miR-199a-5p and miR-199a-3p are highly conserved among several species. The regions highlighted in gray indicate the seed sequence. *D - G*, Expression levels of primary miR-199a-1 (D), primary miR-199a-2 (E), mature miR-199a-5p (F), and mature miR-199a-3p (G) were measured by qRT-PCR in cultures of developing hippocampal neurons at 0, 1, 3, 6, and 10 DIV.

### MeCP2 and miR-199a promote dendritic development in cultured hippocampal neurons

To investigate the influence of the MeCP2/miR-199a axis in the dendritic development of neurons, we expressed MeCP2 or miR-199a in primary cultured hippocampal neurons by infecting with lentiviral systems expressing GFP as control, along with MeCP2 or miR-199a (Fig. 2A). Then, we evaluated dendritic morphology by immunocytochemical assays utilizing antibodies against MAP2, a marker for neuronal dendrites. In neurons expressing exogenous MeCP2, total dendrite length and branch number were found to be significantly higher than that in control. Interestingly, we found that miR-199a expression in neurons significantly increased dendrite length and branch number to the same level as in MeCP2-expressing cells (Fig. 2B-D). We also performed Sholl analysis by measuring the number of dendrites that cross concentric circles at different radial distances from the cell body. The Sholl analysis revealed that miR-199a, as well as MeCP2, enhances dendritic arborization compared to the control (Fig. 2E). Together, these results suggest that the MeCP2/miR-199a pathway promotes dendritic morphogenesis in hippocampal neurons.

**Fig. 2.**
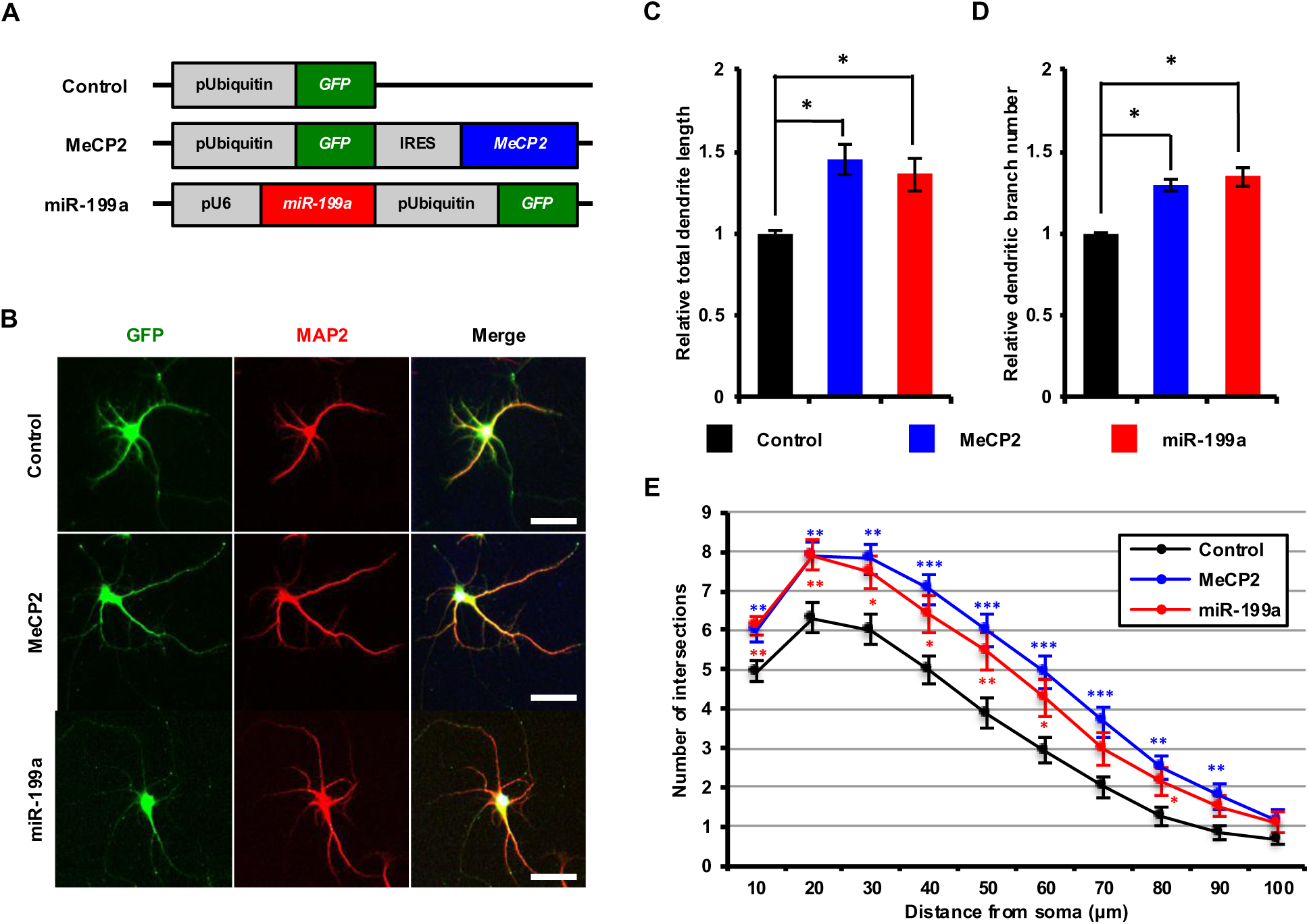
MeCP2 and miR-199a promote dendritic development of cultured hippocampal neurons. *A*, Diagrammatic representations of lentiviral vector constructs. GFP and MeCP2 are expressed under the ubiquitin promoter. miR-199a is expressed as a short hairpin RNA under the U6 RNA polymerase III promoter, and GFP is expressed under the ubiquitin promoter. *B*, Representative images of hippocampal neurons, labeled with anti-MAP2 (red) and anti-GFP (green) antibodies at 6 days *in vitro* (DIV). *C and D*, Quantification of total dendrite length (C) and dendrite branch number (D) in *B*. Student’s *t*-test, *, p < 0.05. n = 4 independent experiments; at least 50 neurons were analyzed in each experiment. *E*, quantification of dendrite complexity by Sholl analysis of neurons infected with lentiviruses co-expressing MeCP2 and miR-199a at 6 DIV. Student’s *t*-test, *, p < 0.05; **, p < 0.01; ***, p < 0.001. Fifty neurons were analyzed at each condition.

### Both mature forms of miR-199a are required for dendritic development in hippocampal neurons

The pre-miR-199a generates 2 types of mature miRNAs, miR-199a-5p and −3p. We, therefore, investigated which of these 2 mature forms contributes to dendritic morphogenesis of neurons. We expressed the miRNA inhibitor (sponge) against either miR-199a-5p (miR-199a5pi) or miR-199a-3p (miR-199a-3pi) in primary cultured hippocampal neurons (Fig. 3A) and evaluated the resultant dendrite morphology. Blocking of endogenous miR-199a-5p was found to lead to decreased total dendrite length, but not branch number, compared with control samples, whereas miR-199a-3p inhibition decreased both (Fig. 3B-D). Furthermore, expression of both miR-199a-5pi and miR-199a-3pi reduced dendritic complexity, although the effect of miR-199a-5pi was less pronounced compared with that of miR-199a-3pi (Fig. 3E). These results suggest that both mature forms of miR-199a can regulate dendritic development, although miR-199a-3p plays a more crucial role than miR-199a-5p.

**Fig. 3.**
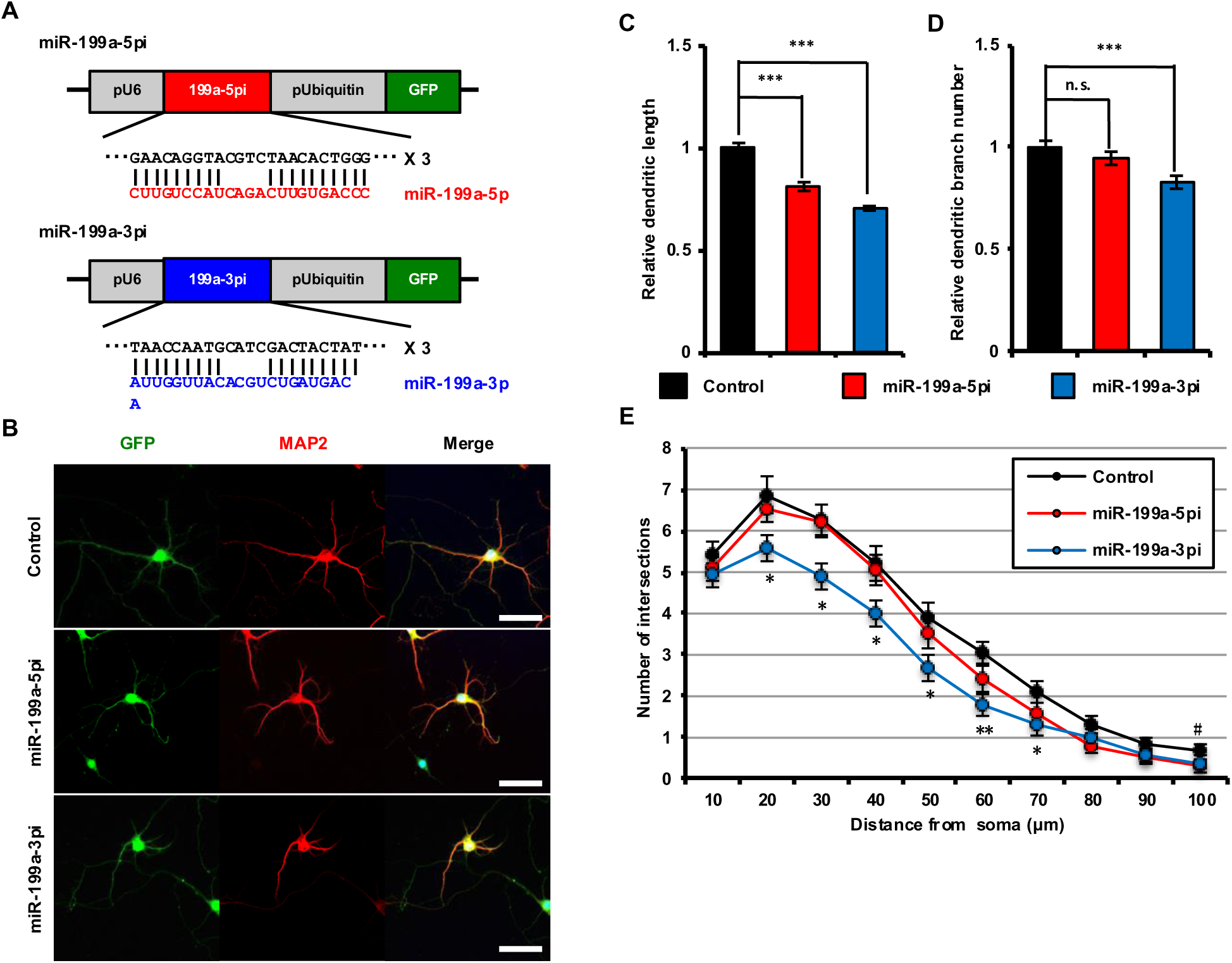
Both mature forms of miR-199a are required for dendritic development in hippocampal neurons. *A*, diagrammatic representations of lentiviral vector constructs expressing sponges against miR-199a-5p and miR-199a-3p. These sponges are expressed as a short hairpin RNA under the U6 RNA polymerase III promoter, and GFP is expressed under the ubiquitin promoter. *B*, representative images of hippocampal neurons infected with lentivirus expressing sponges against miR-199a-5p and miR-199a-3p. These neurons were labeled with anti-MAP2 (red) and anti-GFP (green) antibodies at 6 days *in vitro* (DIV). *C and D*, quantification of total dendrite length (C) and dendrite branch number (D) in *B*. Student’s *t*-test, ***, p < 0.001; n.s., not significant. n = 4 independent experiments; at least 50 neurons were analyzed in each experiment. *E*, quantification of dendrite complexity by Sholl analysis of neurons infected with lentiviruses expressing sponges against miR-199a-5p and miR-199a-3p at 6 DIV. Student’s *t*-test, *, p < 0.05; **, p < 0.01. Fifty neurons were analyzed for each condition.

### miR-199a acts downstream of MeCP2 in the regulation of dendritic development

Since MeCP2 is known to promote the processing of pri-miR-199a, we next examined whether miR-199a functions downstream of MeCP2 in dendrite formation. We expressed the inhibitors for the respective mature miR-199a, along with MeCP2, in cultured hippocampal neurons and evaluated the resultant dendritic morphology (Fig. 4A). Inhibition of miR-199a-5p decreased the MeCP2-mediated enhancement of total dendrite length, but not branch number, in neurons compared with control, whereas blocking miR-199a-3p reduced both (Fig. 4B-D). Moreover, although the effect of miR-199a-3pi was stronger than that of miR-199a-5pi, we found that expression of both the mature forms inhibited MeCP2-induced increase in dendritic arborization (Fig. 4E). These results suggest that both forms of mature miR-199a act as regulatory factors downstream of MeCP2 in dendritic development, although miR-199a-3p is more stringently involved in this regulation.

**Fig. 4.**
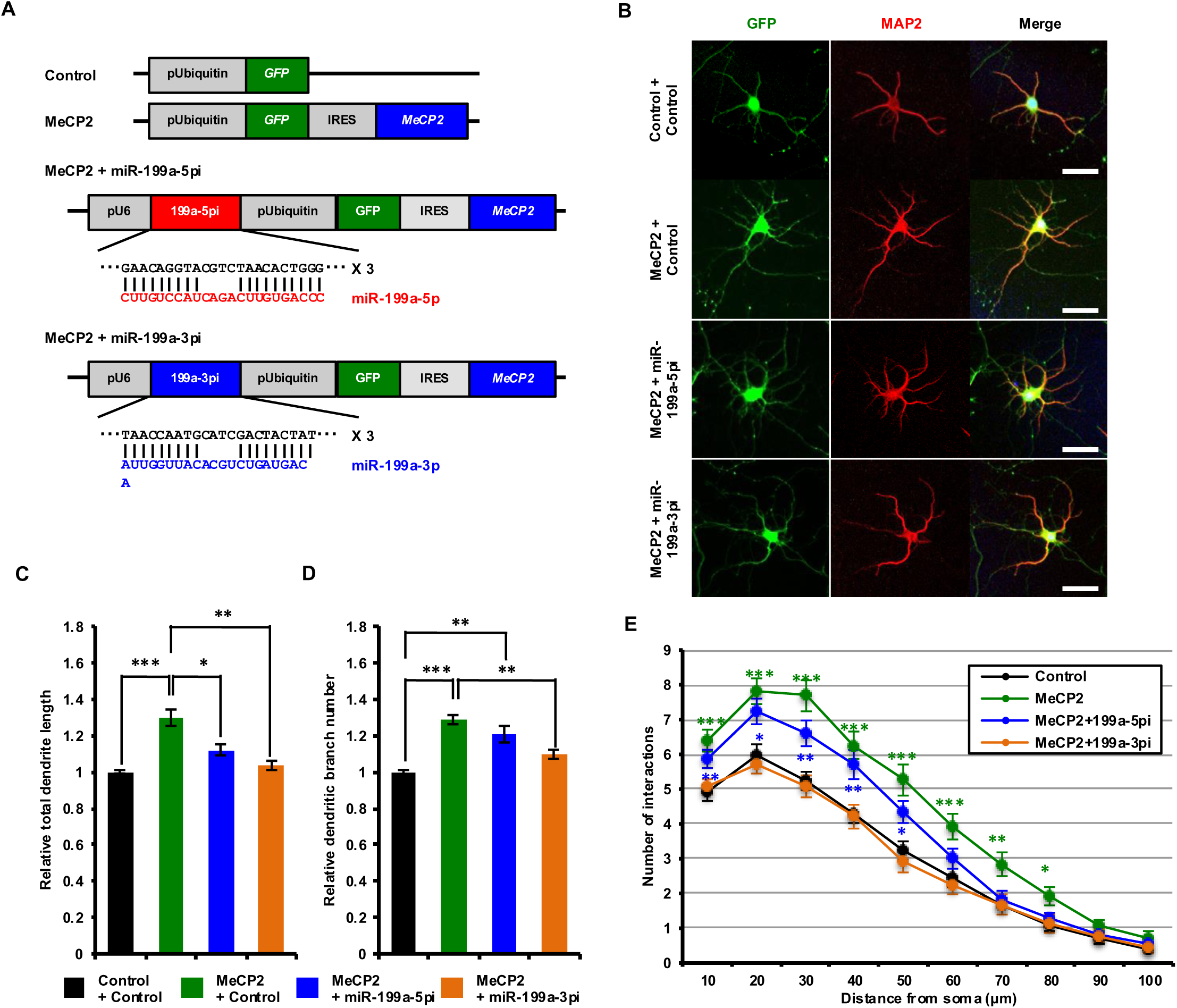
Inhibition of miR-199a counteracts the effect of MeCP2 expression in hippocampal neurons. *A*, diagrammatic representations of lentiviral vector constructs expressing MeCP2 and sponges against miR-199a-5p or miR-199a-3p. These sponges are expressed as a short hairpin under the U6 RNA polymerase III promoter, and GFP and MeCP2 are expressed under the ubiquitin promoter. *B*, representative images of hippocampal neurons infected with lentivirus expressing MeCP2 and sponges against miR-199a-5p or miR-199a-3p. These neurons were labeled with anti-MAP2 (red) and anti-GFP (green) antibodies at 6 DIV. *C and D*, quantification of total dendrite length (C) and dendrite branch number (D) in *B*. Student’s *t*-test, *, p < 0.05; **, p < 0.01; ***, p < 0.001. n = 4 independent experiments; at least 50 neurons were analyzed in each experiment. *E*, quantification of dendrite complexity by Sholl analysis of neurons infected with lentiviruses expressing MeCP2 and sponges against miR-199a-5p or miR-199a-3p at 6 DIV. Student’s *t*-test, *, p < 0.05; **, p < 0.01; ***, p < 0.001. Fifty neurons were analyzed for each condition.

### MeCP2 and miR-199a enhance dendritic development in vivo

We next examined the effects of MeCP2 and miR-199a in dendritic morphogenesis *in vivo* in the mouse brain. For this, we expressed MeCP2 and miR-199a in WT mouse brains by *in utero* electroporation at E14.5 and analyzed neuronal morphology at P10. Almost all the GFP-positive transfected neurons at this stage were located in layer IV (31). When we co-expressed MeCP2 and GFP in the WT mouse brain, electroporated neurons showed enhanced dendrite length and branching number compared with that of control (Fig. 5A-D). Furthermore, co-expression of miR-199a and GFP in these WT neurons significantly promoted increase in dendrite length and branching number, such that the latter were comparable to those in MeCP2-expressing neurons (Fig. 5A-D). We also performed Sholl analysis and found that expression of MeCP2 and miR-199a facilitates dendritic complexity and increases it to levels similar to that in control conditions (Fig. 5E). Taken together, these data suggest that the MeCP2/miR-199a axis positively regulates dendritic development *in vivo*.

**Fig. 5.**
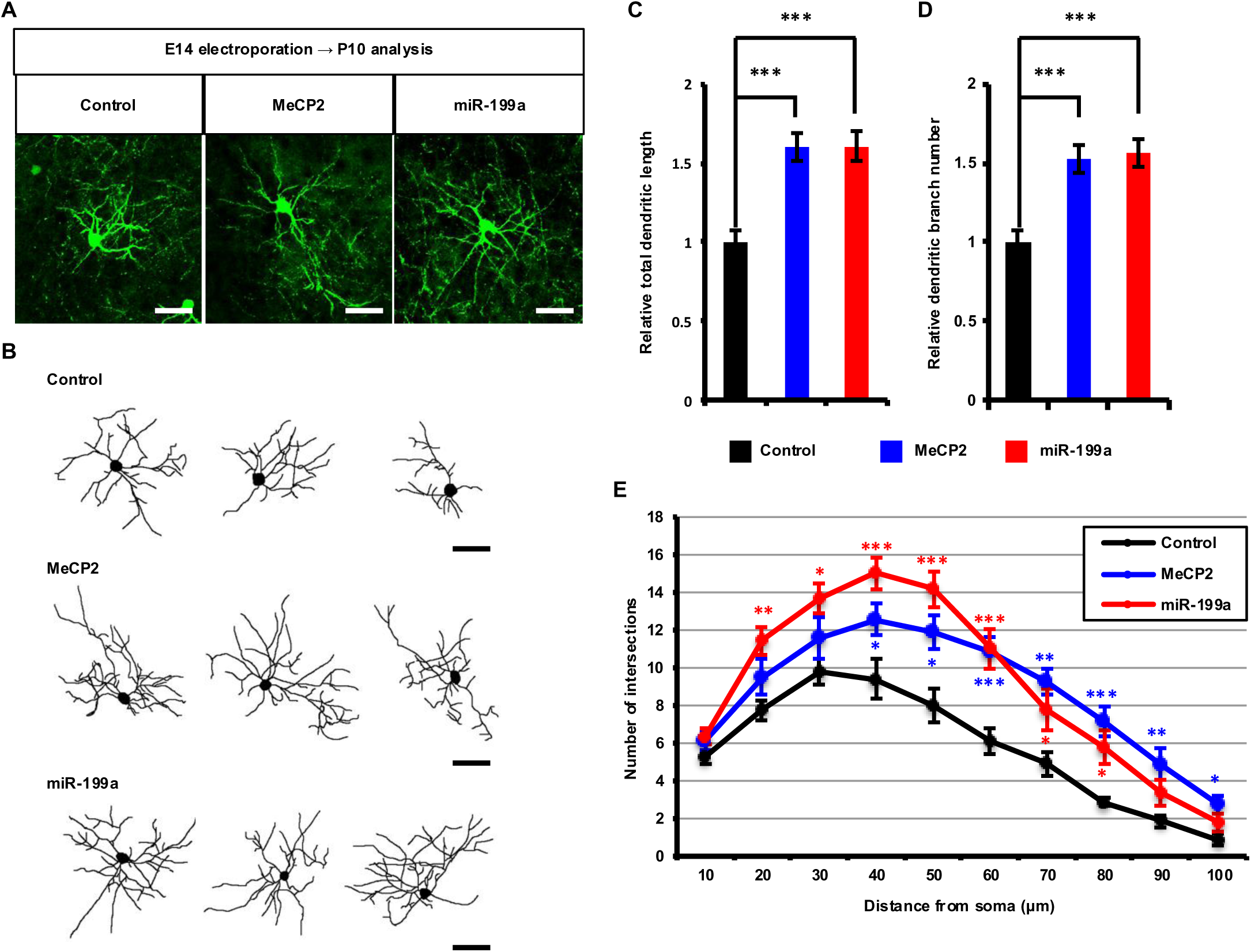
MeCP2 and miR-199a promote dendritic development *in vivo*. *A*, representative image of a transfected brain section labeled with anti-GFP antibody. WT mice embryos were transfected with plasmids expressing only GFP (control), or GFP along with MeCP2 (MeCP2) or miR-199a (miR-199a) by *in utero electroporation* at E14 and sacrificed at P10. Scale bar, 50 µm. *B*, representative tracing images of neurons transfected with the constructs described in *A*. Scale bar, 50 µm. *C and D*, quantification of total dendrite length (C) and dendrite branch number (D) in *B*. One-way ANOVA, Tukey’s post-hoc test, ***, p < 0.001. Number of cells analyzed: control, 13 neurons from three brains; MeCP2, 14 neurons from three brains; miR-199a, 12 neurons from three brains. *E*, quantification of dendrite complexity by Sholl analysis of neurons in *C*. One-way ANOVA, Tukey’s post-test, *, p < 0.05; **, p < 0.01; ***, p < 0.001.

### Expression of miR-199a rescues dendritic impairment in MeCP2-deficient neurons in vivo

We next asked whether miR-199a expression could reverse the impairment of dendritic morphogenesis in MeCP2-deficient neurons. To this end, we expressed miR-199a in the MeCP2-KO mouse brain by *in utero* electroporation at E14.5 and analyzed neuronal morphology at P10. When we expressed only GFP in the MeCP2-KO mouse brain, electroporated neurons showed decreased dendrite length and branching number compared with that in WT. By contrast, the expression of miR-199a in MeCP2-KO neurons significantly reversed the reduced dendrite length and branching number to the level of WT neurons (Fig. 6A-D). Furthermore, Sholl analysis revealed that miR-199a expression restored the dendritic complexity to levels observed in MeCP2-KO neurons (Fig. 6E). Together with the *in vitro* results, these insights support the notion that miR-199a functions as a downstream regulator of MeCP2 in dendritic development *in vivo*.

**Fig. 6.**
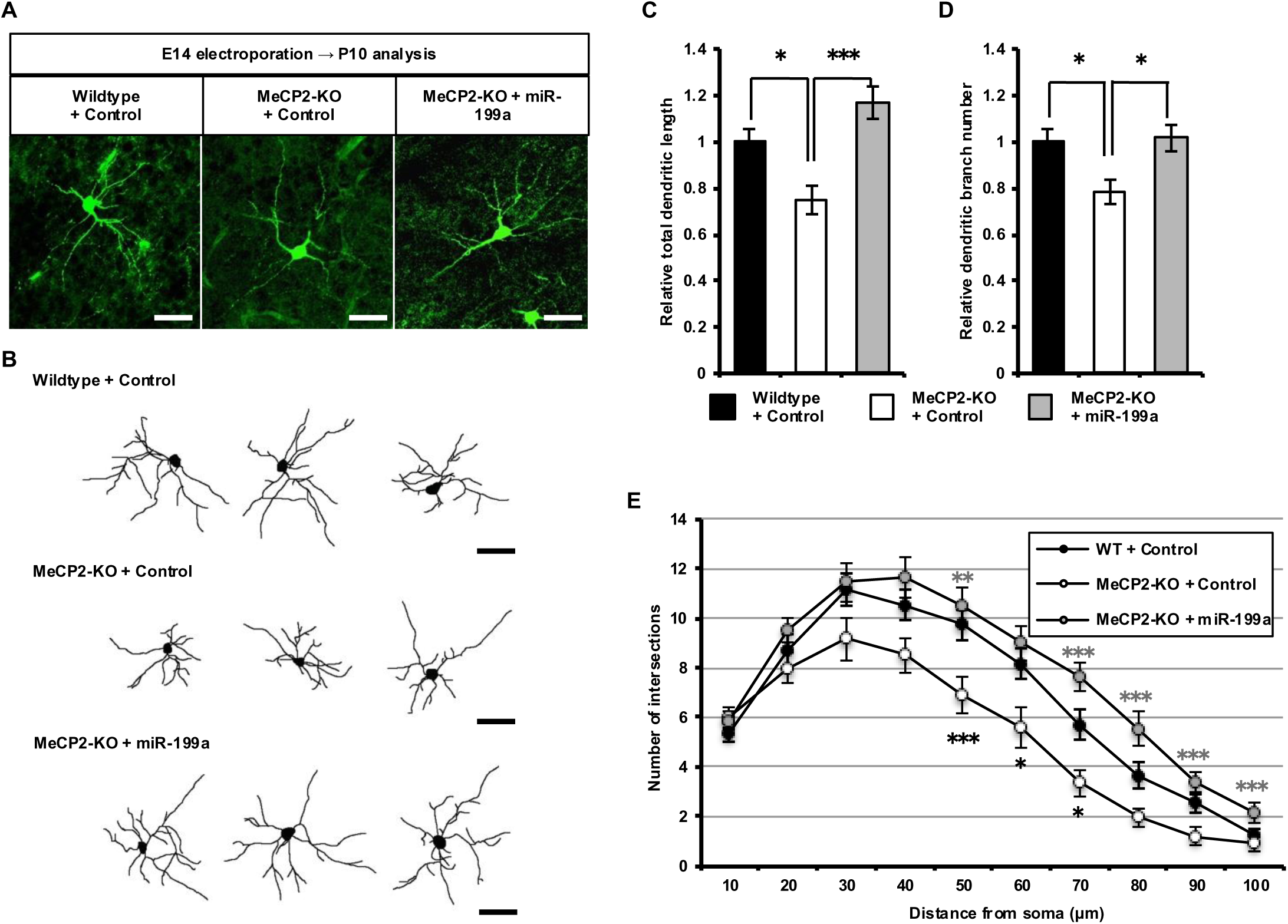
Expression of miR-199a reverses dendritic impairment in MeCP2 knockout neurons *in vivo*. *A*, Representative image of a transfected brain section labeled with anti-GFP antibody. WT and MeCP2-KO mice embryos were transfected with plasmids expressing only GFP (control), or GFP together with miR-199a (miR-199a) by *in utero electroporation* at E14 and sacrificed at P10. Scale bar, 50 µm. *B*, Representative tracing images of neurons transfected with the constructs described in *A*. Scale bar, 50 µm. *C and D*, Quantification of total dendrite length (C) and dendrite branch number (D) in *B*. One-way ANOVA, Tukey’s post-test, *, p < 0.05; ***, p < 0.001. Number of cells analyzed: WT + control, 19 neurons from three brains; MeCP2-KO + control, 17 neurons from three brains; MeCP2-KO + miR-199a, 16 neurons from three brains. *E*, Quantification of dendrite complexity by Sholl analysis of neurons in *C*. One-way ANOVA, Tukey’s post-test, *, p < 0.05; **, p < 0.01; ***, p < 0.001.

### miR-199a-3p targets 3’UTR of Qki mRNA and suppresses its protein expression in neurons

To understand the mechanisms by which miR-199a regulates dendritic morphogenesis downstream of MeCP2, we attempted to identify genes targeted by each mature form of miR-199a. We have previously demonstrated that miR-199a-5p positively regulates mTOR signaling by targeting *Pde4d*, *Sirt1*, and *Hif1a* in hippocampal neurons (32). Since mTOR signaling is known to promote morphological development of dendrites (36), we reasoned that miR-199a-5p enhances dendritic arborization through mTOR signaling. Additionally, since we have earlier shown that miR-199a-3p regulates dendritic development to a greater extent compared with miR-199a-5p, we particularly focused on identifying the genes targeted by miR-199a-3p. To this end, we first utilized public databases such as miRDB (http://mirdb.org), TargetScan (http://www.targetscan.org/vert_72/), and PicTar (https://pictar.mdc-berlin.de), and identified 9 candidate target genes which were common in results from the three databases. These included *amyloid beta (A4) precursor-like protein 2 (APLP2), Cbp/p300-interacting transactivator, with Glu/Asp-rich carboxy-terminal domain, 2 (Cited2), F-box and WD-40 domain protein 11 (Fbxw11), far upstream element (FUSE) binding protein 1 (Fubp1), mitogen-activated protein kinase kinase kinase 4 (Map3k4), muscleblind-like 1 (Mbnl1), platelet-derived growth factor receptor (Pdgfr), protein phosphatase 2 regulatory subunit B’ epsilon (Ppp2r5e), and quaking (Qki)* (Fig. 7A). Since the 3’UTR of *Qki* mRNA has the largest number of seed sites among these candidates and we have recently shown that Qki negatively regulates dendritic development (31), we chose the *Qki* gene as the most plausible target of miR-199a-3p. Next, to investigate whether miR-199a-3p directly targets the *Qki* mRNA, we performed a luciferase reporter assay using reporter constructs in which a luciferase reporter gene was placed under post-transcriptional control of the native *Qki* 3’UTR or a *Qki* 3’UTR harboring mutations in all three miR-199a-3p seed sites (Fig. 7B). These luciferase constructs, along with a miR-199a-expressing plasmid, were co-transfected in hippocampal neurons, and luciferase activity was measured. The latter showed significant decrease on miR-199a expression in neurons containing the construct harboring all three native seed sequences. However, the luciferase activities of reporter constructs with mutation in either of the three seed sequences did not show such sensitivity to miR-199a expression (Fig. 7C). To determine whether miR-199a-3p regulates Qki expression at the protein level, we expressed miR-199a in hippocampal neurons and carried out immunoblotting assays using an anti-Qki antibody. The *Qki* gene is known to generate three isoforms of the Qki protein (Qki-5, Qki-6, and Qki-7) by alternative splicing (37). We found that the expression levels of all three Qki isoforms were substantially decreased in miR-199a-expressing cells compared with that in control cells (Fig. 7D and E). These findings suggest that miR-199a-3p suppresses Qki protein expression in hippocampal neurons through interaction with the 3’UTR of *Qki* mRNA.

**Fig. 7.**
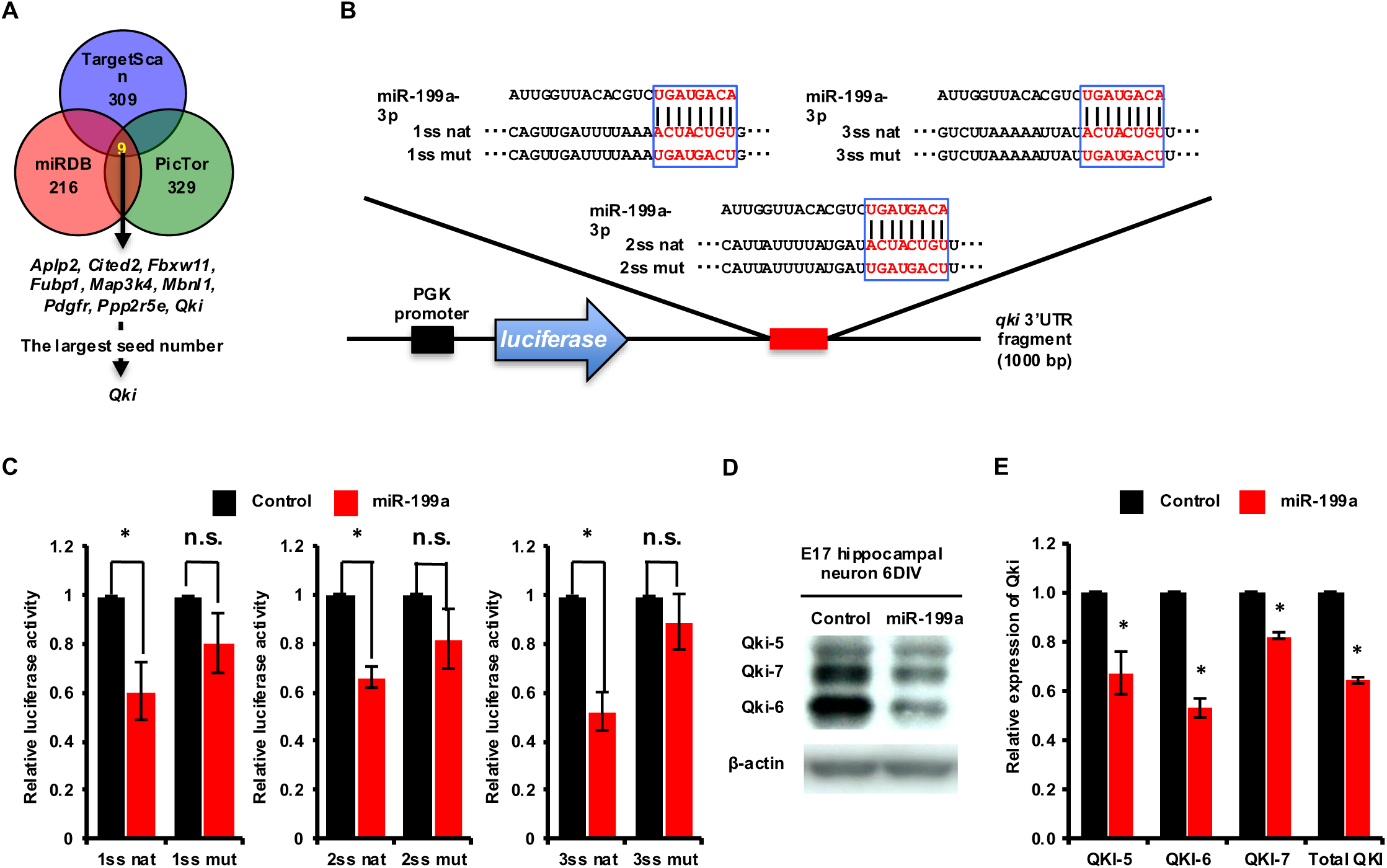
miR-199a-3p targets *Qki* in hippocampal neurons. *A*, Flow chart showing the screening process for selection of downstream targets of miR-199a-3p. *B*, Diagrammatic representations of luciferase reporter constructs that were inserted in a part of the Qki 3’UTR DNA fragment including each seed site, showing the miR-199a-3p sequence, and each miR-199a-3p target sequence of native type (nat) and its mutated sequence (mut). The 5’ seed sequences of miR-199a-3p are marked by blue boxes. *C*, Luciferase activity of *Qki* 3’UTR was suppressed by miR-199a overexpression at 3 days after transfection (left). Student’s *t*-test, *, p < 0.05; n.s., not significant. n = 3 independent experiments. *D*, Western blotting analysis showing that the levels of all three isoforms of Qki protein were decreased in hippocampal neurons infected with lentivirus vector constructs expressing miR-199a at 6 days *in vitro* (DIV). *E*, Quantification of total Qki protein level in *B*. Student’s *t*-test, *, p < 0.05. n = 3 independent experiments.

### miR-199a-2-KO neurons show reduced dendritic formation and Qki knockdown reinstates abnormal dendritic morphology in miR-199a-2-deficient neurons

We have previously generated miR-199a-2 deficient (miR-199a-2-KO) mice, which recapitulates many RTT phenotypes such as reduction in body weight, brain size, and neuronal soma size (32). We, therefore, attempted to confirm that miR-199a-3p regulates Qki expression using the miR-199a-2-KO mouse model by estimating the expression levels of miR-199a and Qki in miR-199a-2-KO neurons. Although miR-199a is supposed to be expressed from 2 gene loci, expression levels of miR-199a-5p and miR-199a-3p in the cortex of miR-199a-2 KO mice were significantly reduced compared with those in WT mice (Fig. 8A). On the contrary, the expression of Qki protein was increased in the same mice (Fig. 8B and C). Then, we investigated the dendritic morphology of miR-199a-2 KO neurons and found that cultured hippocampal neurons from miR-199a-2 KO mice exhibits decreased dendrite length and branching number compared with neurons from WT mice. Further, we investigated whether downregulation of Qki expression could reverse the dendritic phenotypes of miR-199a-2 KO neurons. For this, we transfected miR-199a-2 KO neurons with lentivirus expressing shRNAs against *Qki* and evaluated the resultant dendritic morphology. Interestingly, expression of shRNA against *Qki* in miR-199a-2 KO neurons significantly restored the decrease in dendrite length and branch number (Fig. 8D-F). Decreased dendritic complexity of miR-199a-2 KO neurons was also improved by Qki knockdown (Fig. 8G). From these results, we concluded that miR-199a controls dendrite morphogenesis by targeting Qki.

**Fig. 8.**
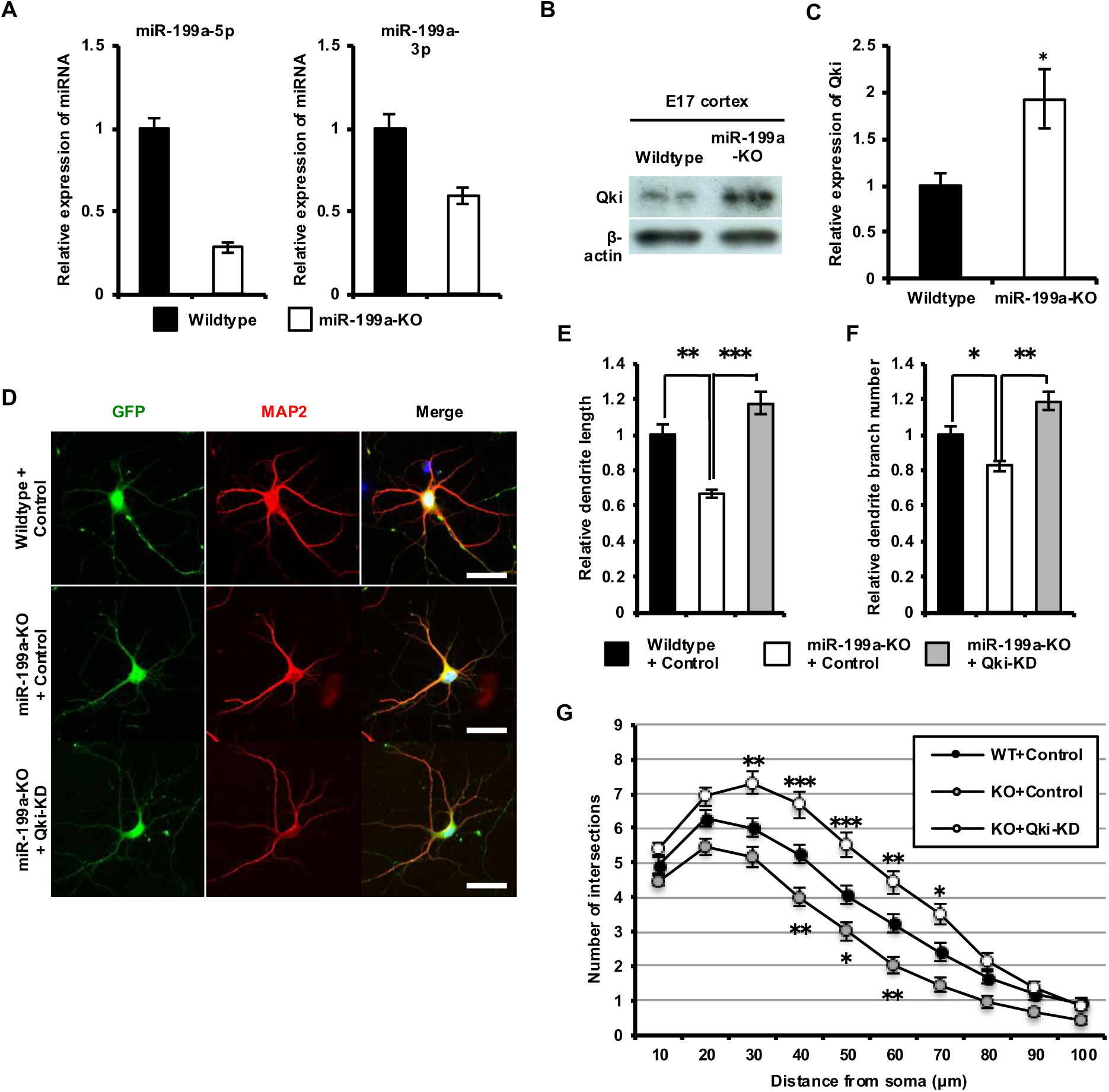
miR-199a-deficient neurons show reduced dendrite formation and Qki knockdown restores abnormal dendritic phenotype. *A*, Expression levels of mature forms of miR-199a-5p and miR-199a-3p in miR-199a-2-KO cortex at E17 by qRT-PCR assays. *B*, Representative immunoblots of Qki, generated by immunoblotting with anti-Qki antibody in miR-199a-2-KO mice cortex for estimation of Qki protein levels. *C*, Quantification of Qki protein levels in *B*. Student’s *t*-test, *, p < 0.05. n = 3 independent experiments. *D*, Representative images of cultured hippocampal neurons from miR-199a-2-KO mice as indicated, infected with lentivirus vector constructs expressing shRNA against *Qki* and labeled with anti-GFP (green) and anti-MAP2 (red) antibodies at 6 days *in vitro* (DIV). *E and F*, Quantification of total dendrite length (E) and dendrite branch number (F) in *D*. Student’s *t*-test, *, p < 0.05; **, p < 0.01; ***, p < 0.001. n = 4 independent experiments; at least 50 neurons were analyzed in each experiment. Scale bar, 50 µm. *G*, Quantification of dendrite complexity by Sholl analysis of neurons from miR-199a-2-KO mice infected with lentivirus vector constructs expressing shRNA against *Qki*. Student’s *t*-test, *, p < 0.05; **, p < 0.01; ***, p < 0.001. Fifty neurons were analyzed for each condition.

## Discussion

Although Rett syndrome is known to be caused by loss-of-function mutations in the *MECP2* gene, the mechanisms by which dysfunction of *MECP2* leads to this disease phenotype has remained largely elusive. A number of studies have reported abnormal reductions in dendritic development in the postmortem brains of RTT individuals and genetically engineered mouse models (9–11, 14, 17, 18, 34, 35). Additionally, MeCP2-deficient neurons are known to exhibit abnormally decreased dendritic development in *in vitro* culture systems (19, 20). Dendritic atrophy found in the brain of RTT patients and mouse models of the disease have been related to dysfunctions of the neural circuits (3). Therefore, these findings imply that defects in dendritic morphology are a key feature in understanding RTT pathogenesis. However, the molecular basis linking MeCP2 functions to the regulation of dendritic development is still unclear. In this study, we have demonstrated the functional role of miR-199a in the MeCP2-mediated regulation of dendritic morphogenesis. We first showed that the expression of MeCP2 correlates with that of miR-199a in developing hippocampal neurons. Transient up-regulation of pri-miR-199a expression was observed at early developmental stages of these cultured hippocampal neurons, along with a reduction in its expression at later stages. On the contrary, the expression levels of both MeCP2 and mature miR-199a were upregulated during maturation of hippocampal neurons. These observations led us to hypothesize that miR-199a has a critical role downstream of MeCP2 in dendritic maturation and implied that the expression of mature miR-199a is tightly regulated at the posttranscriptional level during neuronal maturation. Since we have previously shown that miR-199a regulates multiple aspects of neurons, such as neuronal growth and excitatory synaptic transmission and formation downstream of MeCP2 (32), we postulated that miR-199a could also contribute to dendritic development downstream of MeCP2. In support of our idea, we showed here that miR-199a positively regulates morphological development of neuronal dendrites. Additionally, blocking of mature miR-199a function was shown to abolish the MeCP2-mediated increase in dendritic outgrowth. The expression of miR-199a in MeCP2-KO neurons was found to improve the loss in dendritic morphology elicited by MeCP2 deficiency. In the present study, we also investigated dendritic morphology of miR-199a-2 KO neurons and found that dendrite formation was significantly decreased compared with WT, similar to that observed in MeCP2 KO neurons. Since miR-199a-2 KO mice mimics various phenotypes of MeCP2 KO mice (32), miR-199a KO neurons understandably recapitulated the dendritic phenotype of MeCP2 KO neurons. Taken together, these findings clearly indicate that miR-199a acts downstream of MeCP2 in the regulation of dendrite morphogenesis and support the notion that miR-199a is a critical downstream target of MeCP2 in RTT pathogenesis (32).

We also highlighted the differential effects of mature miR-199a, miR-199a-5p, and mir-199a-3p on dendritic development. Whereas inhibition of miR-199a-5p reduced only dendrite length, blocking miR-199a-3p induced a decrease in both dendrite length and branch number. These results suggested that both mature forms of miR-199a regulate dendrite formation through independent cascades by targeting different downstream targets. We have earlier demonstrated that miR-199a-5p positively regulates neuronal functions including neuronal growth, excitatory synaptic transmission, and synaptogenesis by controlling mTOR signaling, and mTOR signaling has been reported to enhance dendrite outgrowth (32, 36). Based on this, we postulated that miR-199a-5p promotes dendritic formation, at least in part, by regulating mTOR signaling. On the contrary, the target genes of miR-199a-3p in neurons have not been investigated so far. Since *Qki* mRNA has three seed sequences of miR-199a-3p in its 3’UTR and we recently reported the negative regulation of dendrite development by Qki (31), we hypothesized that miR-199a-3p targets *Qki* gene in neurons. Our findings were in overall agreement with the hypothesis and demonstrated that miR-199a-3p directly targets the 3’UTR of *Qki* mRNA and suppresses Qki protein expression in neurons.

In the present study, the mechanisms underlying Qki-mediated regulation of dendritic development remain to be determined. Qki is an RNA-binding protein that belongs to the signal transduction and activation of RNA (STAR) protein family and is reported to regulate RNA splicing, RNA export, RNA stability, and protein translation (38, 39). A previous study has reported that Qki regulates the stability of actin-interacting protein 1 (AIP1) mRNA during oligodendrocyte differentiation (40). Parallelly, a number of studies have revealed that actin cytoskeleton dynamics plays an important role in the regulation of dendritic morphogenesis (41, 42). Therefore, it is possible that Qki regulates dendritic formation by targeting AIP1. Many other target mRNA candidates of Qki have also been identified by SELEX (systematic evolution of ligands by exponential enrichment) and bioinformatics based analysis (43). Intriguingly, *Tbr1*, *Wnt2a*, and *Rab5a* have been identified as putative targets of Qki. A recent work has reported that Tbr1 regulates laminar patterning of retinal ganglion cell dendrites (44). Another study has shown that the Wnt/Dishevelled pathway controls development of dendritic morphology and a component of the early endocytic pathway, Rab5a, enhances dendritic morphogenesis of fly dendritic arborization neurons (45).

Genetic studies have implicated *Qki* in psychiatric disorders including schizophrenia (46, 47), and mutations in *MECP2* have been reported in schizophrenia patients (48). Given the dual lines of evidence on the contribution of altered dendritic morphology in various neurological diseases (5, 49, 50) and MeCP2-mediated regulation of Qki expression, it is possible that dysfunction of MeCP2/miR-199a/Qki axis contributes to abnormal dendritic phenotypes associated with a variety of neurological disorders.

Although an abnormal decrease in dendritic development is observed in the postmortem brains of RTT patients and mouse models, the mechanisms underlying this dendritic phenotype have not been understood to date. Our findings reveal a key role of the MeCP2/miR-199a/Qki pathway in the regulation of dendritic morphogenesis and provide a molecular basis linking MeCP2 dysfunction with impaired dendritic development. We, therefore, propose that controlling this molecular pathway offers a new avenue for the development of an effective therapeutic strategy for RTT.

## Material and Methods

### Animals

All aspects of animal care and treatment were carried out according to the guidelines of the Experimental Animal Care and Use Committee of Kyushu University and Nagoya University, respectively. Timed-pregnant ICR mice (Japan SLC, Shizuoka, Japan) were used for the present research. The *Mecp2* mutant (*Mecp2^tm1.1Jae^*) used in this study was created by deleting exon 3 containing the methyl-DNA-binding domain of MeCP2 (12) and was obtained from Jackson Laboratories. miR-199a-2-KO mice (Acc. No.: CDB1185K; http://www2.clst.riken.jp/arg/micelist.html) were generated as described previously (32). All mutant mice were maintained on a C57BL/6J (Jackson Laboratories) or ICR (Japan SLC) background.

### Cell culture

Hippocampal neuronal cultures were carried out as previously described (51). Briefly, hippocampi were dissected from embryonic day 17.5 (E17.5) mice and were digested in S-MEM (Gibco) containing 0.1% papain (Sigma), and triturated with 60 mg/ml DNase I (Sigma) and 10% fetal bovine serum (FBS). Dissociated cells were plated on poly-*L*-lysine (10 µg/ml, Sigma)-coated culture dishes at densities of 1 × 10^4^ cells/cm^2^ and 5 × 10^4^ cells/cm^2^ for immunocytochemical and immunoblotting assays, respectively. Once neurons had adhered (after 3-4 h), the medium was replaced with serum-free neuronal maintenance medium, comprising Neurobasal medium (Invitrogen) supplemented with B-27 (Invitrogen) and 0.5 mM GlutaMAX (Invitrogen). Cytosine β-D-arabinofuranoside hydrochloride (Sigma) was added 1 day after plating to prevent the proliferation of undifferentiated and glial cells. The medium was changed every 3 days. HEK 293T cells were cultured in DMEM containing 10% FBS and gentamicin (Nacalai Tesque).

### Construction of vectors

Lentiviral vectors used to express the short hairpin RNA (shRNA) for *Qki* (pLLX-shQki) and FLAG-tagged Qki (pLEMPRA-Qki and pLEMPRA-Qki-3’UTR) were generously provided by Drs. Z. Zhou and M. E. Greenberg. pLLX and pLEMPRA are dual-promoter lentivirus vectors constructed by inserting the U6 promoter-driven shRNA cassette 5’ into the ubiquitin-C promoter in the FUIGW plasmid (52, 53). The pLLX-primary-miR-199a plasmid was constructed by inserting a PCR-amplified fragment from mouse genomic DNA into the HpaI and XhoI restriction sites of pLLX. The shRNA for Qki and sponges against miR-199a-5p- or miR-199a-3p-expressing lentivirus plasmids were constructed by inserting the following oligonucleotides into the HpaI and XhoI sites of pLLX: sh-Qki-3-Fw, 5’-TGGACTTACAGCTAAACAACTTTTCAAGAGAAAGTTGTTTAGCTGTAAGTCCTTTTTTTGAAC-3’, and sh-Qki-3-Rv, 5’-tcgagttccaaaaaaGGACTTACAGCTAAACAACTTt-ctcttgaaAAGTTGTTTAGCTGTAAGTCCa-3’; sponge-miR-199a-5p-Fw, 5’-gacgttaacGAACAGGTACGTCTAACACTGGGGAACAGGTACGTCTAACACTGGGGAACAGGTACGTCTAACACTGGGttttttctcgaggtc, and sponge-miR-199a–5p-Rv; 5’-gacctcgagaaaaaaCCCAGTGTTAGACGTACCTGTTCCCCAGTGTTAGACGTACCTGTTCCCCAGTGTTAGACGTACCTGTTCgttaacgtc-3’; sponge-miR-199a-3p-Fw, and 5’-gacgttaacTAACCAATGCATCGACTACTATTAACCAATGCATCGACTACTATTAACCAATGCATCGACTACTATttttttctcgaggtc-3’; and sponge-miR-199a-3p-Rv and 5’-gacctcgagaaaaaaATAGTAGTCGATGCATTGGTTAATAGTAGTCGATGCATTGGTTAATAGTAGTCGATGCATTGGTTAgttaacgtC-3’. *Qki*, cDNA fragments and its 3’UTR-containing fragments were amplified by PCR using KOD polymerase (Toyobo) and subsequently cloned into the EcoRI and AscI sites of pLEMPRA-MeCP2 (53). The luciferase reporter plasmids pmirGLO-Qki-3’UTR-1ss-nat, −2ss-nat, and −3ss-nat were constructed by inserting the genomic DNA fragments of Qki 3’UTR (1ss, +1361 to +2374; 2ss, +3330 to +4337; and 3ss, +4134 to +5144) into the PmeI and XhoI sites of pmir-GLO (Promega). Each of the mutated luciferase reporter constructs was obtained by inserting the mutated 3’UTR of Qki with the seed regions of miR-199a-3p into the PmeI and XhoI sites of pmir-GLO.

### In utero electroporation

To investigate dendritic growth *in vivo*, *in utero* electroporation was performed on WT or MeCP2-KO E14 mice (ICR background) embryos, as described previously (54). Briefly, plasmid DNA (0.1 μg/μL in phosphate-buffered saline (PBS) containing 0.1% Fast Green FCF dye) was injected (0.5-1 μl) into the lateral ventricle of the embryonic brain from outside the uterus with a glass micropipette (GD-1, Narishige). While holding the embryo in the uterus with forceps-type electrodes (Nepa Gene), 50-ms electric pulses of 45 V each were delivered 5 times at intervals of 950 ms using a CUY21 Single Cell Electroporator (Nepa Gene). Glass micropipettes were prepared using a P-1000IVF micropipette puller (Sutter). Animals were perfused with 4% paraformaldehyde at postnatal day 10 (P10). Collected brains were postfixed with 4% paraformaldehyde overnight at 4°C and then equilibrated in 30% sucrose. Brains were frozen at −80°C after embedding in Tissue-Tek optimal cutting temperature (OCT) compound (Sakura Finetek) and serially sectioned at 40-μm thickness.

### Immunocytochemical assays

Cells were fixed at the indicated day(s) *in vitro* (DIV) with 4% paraformaldehyde in PBS, washed with PBS, permeabilized, and blocked with blocking buffer (3% FBS and 0.1% Triton X-100 in PBS) at room temperature (RT). The cells were then incubated with the primary antibody solution at RT for 3 h. The cells were then washed with PBS and further incubated with the secondary antibody solution at RT for 1 h. After further washing with PBS, these were mounted on glass slides.

### Immunohistochemical assays

Brain sections were washed with PBS, permeabilized, and blocked with blocking buffer at RT. These sections were then incubated with the primary antibody solution overnight at 4°C, washed with PBS, and further incubated with the secondary antibody solution at RT for 2 h. These sections were mounted on glass slides, after washing with PBS. Fluorescence images were acquired using a Zeiss LSM 700 confocal microscope with a 20× objective lens. Z series of 20 images were taken at 1-µm intervals at a resolution of 1,024 × 1,024 pixels.

### Immunoblotting

Cells were lysed with a buffer containing 0.5% Nonidet P-40, 10 mM Tris-HCl, pH 7.5, 150 mM NaCl, and 1% protease inhibitor mixture (Nacalai Tesque). Lysates were sonicated and centrifuged at 20,000x*g* for 15 min at 4°C. Total cell lysates were subjected to SDS-PAGE and transferred to a PVDF transfer membrane (GE Healthcare). The blots were blocked with 0.3% skim milk in TBST (50 mM Tris-HCl, pH 7.5, 150 mM NaCl, 0.1% Tween 20), incubated with a primary antibody, and washed again with TBST. This was followed by incubation with a peroxidase-conjugated secondary antibody. Immunoreactive bands were detected by enhanced chemiluminescence using the ECL Prime Western Blotting Detection Reagent (GE Healthcare).

### Antibodies

For immunocytochemical and immunohistochemical assays, the following primary antibodies were used: chicken anti-green fluorescent protein (GFP) (1:1000; Aves Labs) and guinea pig anti-microtubule-associated protein 2 (MAP2) (1:1000; Synaptic Systems) antibodies. The following secondary antibodies were used: CF488A donkey anti-chicken IgY (IgG) (H + L) highly cross-adsorbed (1:500; Biotium) and CF555 donkey anti-guinea pig IgG (H + L) highly cross-adsorbed (1:500; Biotium). Nuclei were stained using bisbenzimide H 33258 fluorochrome trihydrochloride (Nacalai Tesque).

For immunoblotting, the following primary antibodies were used: rabbit anti-MeCP2 (1:500; Diagenode), rabbit anti-QKI (1:250; ProteinTech), rabbit anti-GAPDH (1:1000; Cell Signaling Technology), and mouse anti-β-actin (1:1000; Cell Signaling Technology). The following secondary antibodies were used: anti-mouse IgG, HRP-linked whole antibody from sheep (1:5000; GE Healthcare) and anti-rabbit IgG, HRP-linked whole antibody from donkey (1:5000; GE Healthcare).

### Luciferase reporter assay

Hippocampal neurons were co-transfected with control (pLLX) or miR-199a expression vectors and pmir-GLO-Qki-1ss-nat, pmir-GLO-Qki-1ss-mut, pmir-GLO-Qki-2ss-nat, pmir-GLO-Qki-2ss-mut pmir-GLO-Qki-3ss-nat, or pmir-GLO-Qki-3ss-mut using Lipofectamine 2000 (Life Technologies). After transfection, the cells were incubated for 3 days, and lysed with a Reporter Lysis Buffer. Luciferase activity of the lysates was measured with the Dual-Glo Luciferase Assay System (Promega) according to the manufacturer’s protocol. Firefly luciferase activities were determined by three independent transfections and normalized by comparison with the Renilla luciferase activities of the internal control.

### Morphological analysis of cultured hippocampal neurons

For analysis of dendrite development *in vitro*, hippocampal neurons were subjected to immunostaining with antibodies against MAP2 and GFP at 6 DIV. Dendrites were defined as MAP2-positive neurites. Sholl analysis and quantification of dendritic length and branch number were performed using the ImageJ software. For Sholl analysis, concentric circles with radii increasing in 10 µm increments were defined from the center of the cell body. The number of MAP2-positive dendrites crossing each circle was then counted.

For *in vivo* dendrite analysis, brain sections were subjected to immunohistochemical labeling with the anti-GFP antibody. Total dendritic length and branch number of GFP-positive neurons in layer IV were measured using ImageJ. For Sholl analysis, concentric circles with radii increasing in 10 μm incrementsQ were defined from the center of the cell body. The number of GFP-positive dendrites crossing each circle was counted. The total number of crossings for each cell was calculated as an objective measurement of total dendritic complexity.

### Quantitative real time-polymerase chain reaction (qRT-PCR) assays

qRT-PCR analysis was performed to measure the expression levels of primary and mature miRNAs, as described previously (55). In brief, total RNA was extracted with TRIzol (Invitrogen) and subjected to reverse transcription with the SuperScript VILO cDNA Synthesis Kit (Invitrogen) according to the manufacturer’s protocol. qRT-PCR was performed with the StepOne Real-Time PCR System (Applied Biosystems). To detect primary and mature miRNAs, a TaqMan MicroRNA Assay kit (Applied Biosystems) was used in accordance with the manufacturer’s protocol. Data analysis was performed using the comparative Ct method. Results were normalized to the *Gapdh* or *U6* snRNA.

### Statistical analysis

The results presented are the average of at least three experiments, each performed in triplicates, with standard errors. Statistical analyses were performed by one-way ANOVA, followed by Tukey’s multiple comparison tests or by Student’s *t*-test, as appropriate, using Prism 5 (GraphPad software). Probabilities of *p* < 0.05 were considered significant.

## Acknowledgments

This work was funded by the Japan Society for the Promotion of Science (JSPS) KAKENHI (24240051, 17H01390, and 16H06279 (PAGS)), and Intramural Research Grants (27-7 and 30-9) for Neurological and Psychiatric Disorders of NCNP to K.N., Grant-in-Aid for Scientific Research on Innovative Areas (16H06527) to K.N., MEXT/JSPS KAKENHI Grant-in Scientific Research on Innovation Areas “Constructive understanding of multi-scale dynamism of neuropsychiatric disorders” (19H05211) to K.T., JSPS KAKENHI (18K06484) to K.T., JSPS KAKENHI (16K18391) to K.T., JSPS KAKENHI (19K16918) to H.N. a grant from Kawano Masanori Memorial Public Interest Incorporated Foundation for Promotion of Pediatrics (29–11) to K.T., and a NPO Rett syndrome support organization grant (2019-01-09) to K.T. and (2019-02-10) to H.N. This research was also supported by the Japan Agency for Medical Research and Development (AMED) under Grant Number JP19ek0109411 to K.T., JP19dm0207075h to N.O.

We thank M.E. Greenberg and Z. Zhou for sharing reagents and all members of the Laboratory of Molecular Neuroscience, Department of Stem Cell Biology and Medicine, Kyushu University and Department of Psychiatry, Nagoya University. We acknowledge technical assistance from the Research Support Center, Research Center for Human Disease Modeling, Kyushu University Graduate School of Medical Science. We also thank Yuko Masuda for technical assistance. We would like to thank Editage (www.editage.com) for English language editing.

## Conflict of Interest Statement

The authors declare no competing financial interests.

## Author Contributions

K.I., K.N., and K.T. designed research; K.I., H.N., M.Y., and K.T. performed experiments; K.I., H.N., F.O., N.O., K.N., and K.T analyzed data; K.I., K.N., and K.T. wrote the manuscript. All authors reviewed the manuscript.

## Notes

### Competing Interest Statement

The authors have declared no competing interest.

### Summary of Updates

Oders of authors updated; author affiliations updated

